# Compulsivity is linked to maladaptive choice variability but unaltered reinforcement learning under uncertainty

**DOI:** 10.1101/2023.01.05.522867

**Authors:** Junseok K. Lee, Marion Rouault, Valentin Wyart

## Abstract

Compulsivity has been associated with variable behavior under uncertainty. However, previous work has not distinguished between two main sources of behavioral variability: the stochastic selection of choice options that do not maximize expected reward (choice variability), and random noise in the reinforcement learning process that updates option values from choice outcomes (learning variability). Here we studied the relation between dimensional compulsivity and behavioral variability, using a computational model which dissociates its two sources. We found that compulsivity is associated with more frequent switches between options, triggered by increased choice variability but no change in learning variability. This effect of compulsivity on the ‘trait’ component of choice variability is observed even in conditions where this source of behavioral variability yields no cognitive benefits. These findings indicate that compulsive individuals make variable and maladaptive choices under uncertainty, but do not hold degraded representations of option values.

---

Compulsivity has been associated with shifts in decision-making under uncertainty^1^. Patients with clinical obsessive-compulsive disorder (OCD) have been shown to need more sensory evidence to commit to a perceptual decision, reflected in increased decision thresholds but no changes in evidence accumulation^2,3^. Similarly, OCD patients show deficits in the reporting but not the learning of uncertain stimulus statistics^4^. These findings have recently been shown to extend to subclinical fluctuations of self-reported ‘compulsivity and intrusive thoughts’ symptoms in the general population^5,6^. OCD and schizophrenia, both of which are characterized by intrusive thoughts, are associated across studies with altered decision confidence^7^. During the learning of uncertain options values through trial-and-error – also known as reinforcement learning (RL), compulsivity has been linked to decision variability, but also the reliance on less complex decision policies^8–11^.

However, these latter findings have been obtained using RL models that assign all decision variability to choice stochasticity. This standard assumption neglects an important source of decision variability: random noise in the RL process that updates option values from choice outcomes^12^. This ‘learning noise’ accounts for a large fraction of decision variability otherwise incorrectly assigned to choice stochasticity^13^. It remains largely unclear whether compulsivity is specifically linked to choice stochasticity, learning noise, or both.

Furthermore, the decision variability observed during reinforcement learning can either be adaptive or maladaptive depending on the volatility of option values^14,15^. When option values change unpredictably over time, variable decisions trigger the exploration of recently unchosen options whose values have become more uncertain^13,16^. By contrast, when option values are stable, variable decisions result in unwarranted choices away from the highest rewarded option. Whether compulsivity is linked to decision variability selectively in volatile environments (where such variability is adaptive) or also in stable environments (where such variability is maladaptive) remains unknown.

Here, we studied human behavior in canonical reinforcement problems (two-armed bandits) involving either stable or volatile option values^17^ (Fig. 1a). We fitted a RL model incorporating both learning noise and choice stochasticity to human decisions to decompose participants’ decision variability into their learning- and choice-based sources^13,17^ (Fig. 2a). We also measured individual differences in a self-reported transdiagnostic dimension of compulsivity in the same participants^6,8^. Across two independent datasets, we obtained replicable evidence that compulsivity is associated with increased choice stochasticity without changes in learning noise. Using a test-retest reliability approach, we further quantified the effect of compulsivity on the ‘trait’ (test-retest reliable) component of choice stochasticity. This effect is present even when option values are stable and choice stochasticity yields no cognitive benefits in terms of exploration.

**Figure 1.**
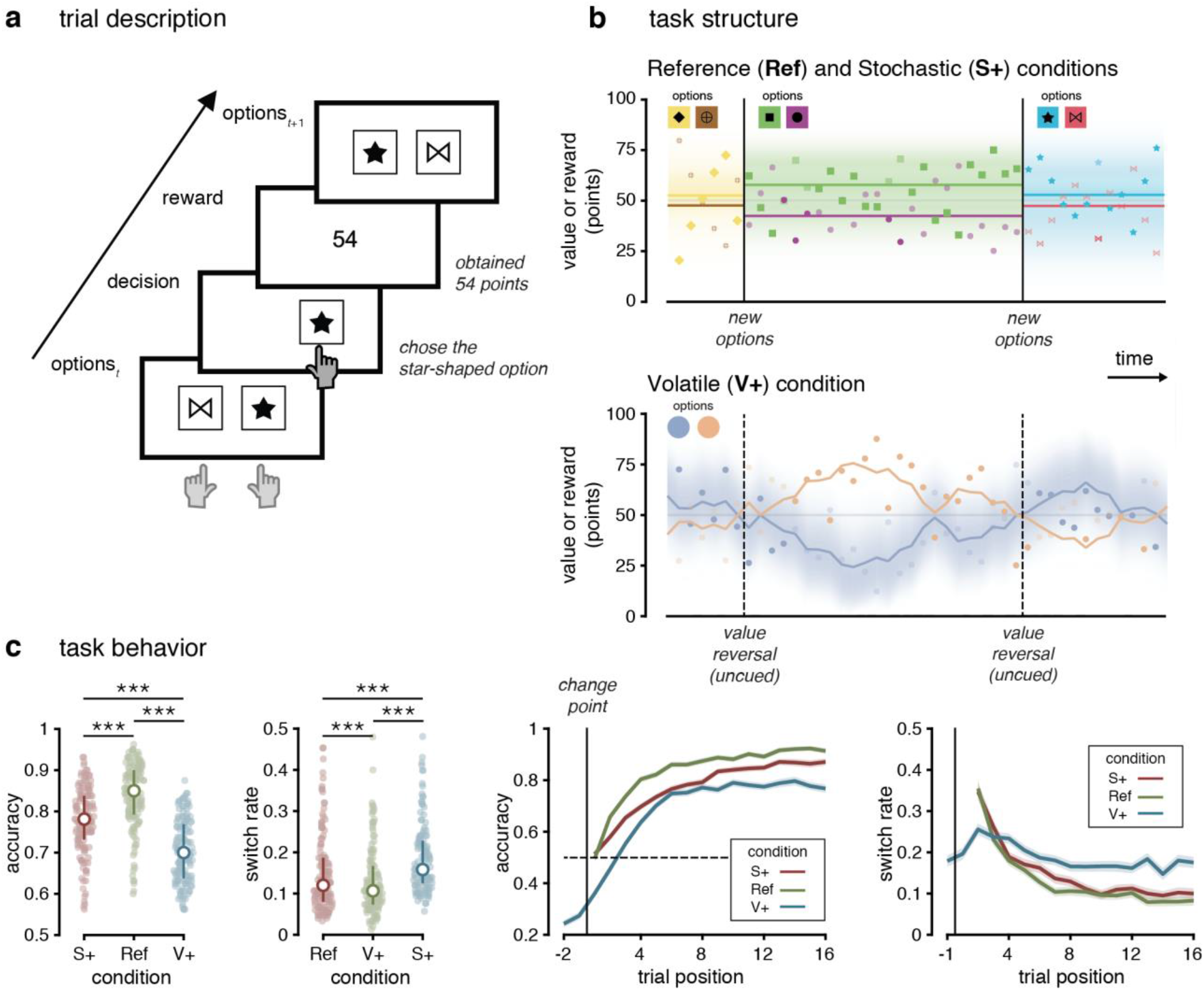
Experimental task and participants’ behavior. (first dataset: *N* = 137). (**a**) Trial description. On each trial, participants were asked to choose between two options. Their chosen option was highlighted, and were then presented with the reward associated with the chosen option in the center of the screen. (**b**) Task structure. Reward schedule of an option during a block. Colored shapes indicate the reward given for an option. The set of shape options are shown for each round. Bold colored shapes indicate the option chosen on a trial. Vertical lines demarcate the explicit beginning and the end of a round in the Reference (Ref) and the Stochastic (S+) conditions and a covert reversal in the Volatile (V+) condition. Upper: Bold colored horizontal lines show the generative mean for the option corresponding to the correct shape in the Reference and Stochastic conditions. Lower: Bold lines indicate the generative reward mean for the best option drifting throughout the course of a block in the Volatile condition. (**c**) Left: Average accuracy and average switch rate in each condition. Colored dots represent individual participants’ mean accuracy and overall switch rate respectively (*N* = 137). White dots indicate the median accuracy and median switch rate respectively. Error bars represent the 1st and 3rd quartiles. Right: Accuracy and proportion of switch decisions as a function of trial position in each condition. Solid lines indicate the mean accuracy and mean switch rate across participants. Shaded areas represent s.e.m. The vertical line represents the start of a new block (Ref, S+) or a reversal (V+).

**Figure 2.**
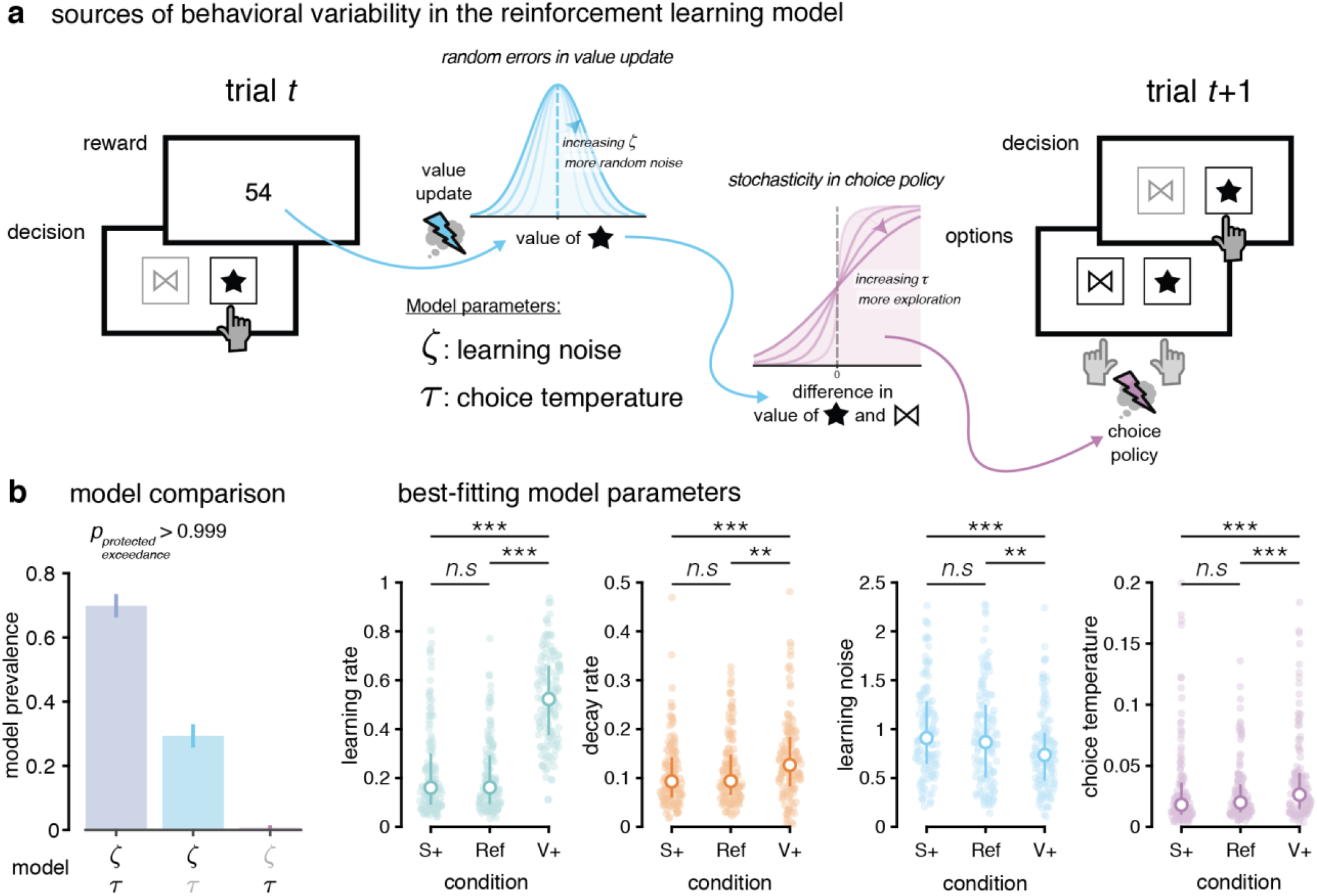
Computational model and participants’ best-fitting parameters. (**a**) Sources of behavioral variability in the reinforcement learning model. Learning noise corrupts the precision with which participants updated the learnt option values. Choice temperature controls the stochasticity of the softmax choice policy. (**b**) Left: Model comparison favors a model with both learning noise and choice stochasticity (left bar), with high prevalence across tested participants. Models lacking either source of decision variability (middle and right bars) provide poorer fits to human behavior. Right: best-fitting individual model parameters in each of the three conditions. White dots indicate the median parameter value and error bars represent the 1^st^ and 3^rd^ quartiles. Circles display individual data points (first dataset, *N* = 137 participants). \*\*\**p*<0.001, \*\**p*<0.01, *n*.*s*.: not significant, Wilcoxon signed-rank tests.

## Results

### Experimental task

We tested the behavior of participants in two-armed bandit problems involving choice options associated with uncertain values^17^ (Fig. 1a; see Methods). In the ‘reference’ (Ref) condition, participants played short rounds of trials with two choice options associated with stable reward distributions. In the ‘stochastic’ (S+) condition, reward distributions had larger variances than the ones used in the reference condition. In the ‘volatile’ (V+) condition, instead of playing short rounds of trials with stable reward distributions, participants played long rounds of trials with choice options associated with drifting reward distributions.

Participants were less accurate at choosing the option associated with the highest reward mean in the S+ and V+ conditions compared to the Ref condition (Fig. 1b; signed-rank test, S+ vs Ref: *z* = − 6.5, *p* < 0.001; V+ vs Ref: *z* = −9.5, *p* < 0.001). Participants also switched more between choice options in these two conditions (S+ vs Ref: *z* = 3.8, *p* < 0.001; V+ vs Ref: *z* = 7.6, *p* < 0.001), and particularly so in the volatile condition (V+ vs S+: *z* = 5.1, *p* < 0.001). These differences between conditions were replicated in the second dataset (Supplementary Fig. 1a).

### Computational model of decision variability

We compared human behavior to a reward-maximizing agent (see Methods). Human behavior was substantially less accurate and more variable than the reward-maximizing agent, in all conditions (Supplementary Fig. 2). To characterize this variability, we fitted a RL model^13,17^ decomposing variability into two sources (Fig. 2a): 1. variability arising from random noise in the update of option values from choice outcomes (learning noise), and 2. variability arising from the stochastic selection of choice options which do not maximize value (choice temperature). We validated this model for the characterization of variability through ‘simulation-recovery’ analyses (see Methods, Supplementary Fig. 3).

**Figure 3.**
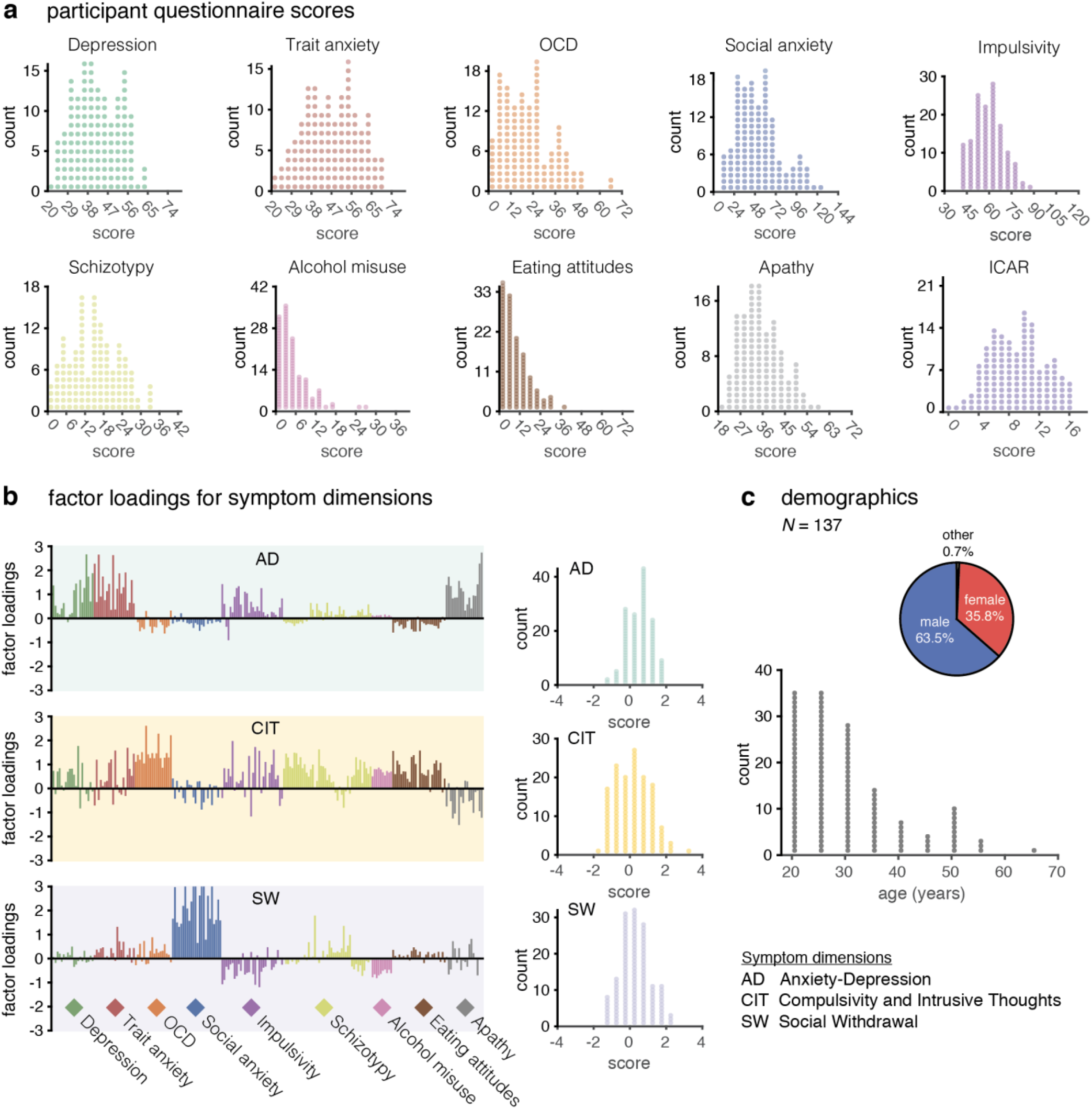
Inter-individual variability in symptom dimension scores and demographics variables. (first dataset, *N* = 137 participants). (**a**) Histograms showing distributions across questionnaire scores in our sample. We evaluated depression using the Self-Rating Depression Scale (SDS), trait anxiety questionnaire using the State Trait Anxiety Inventory (STAI), obsessive-compulsive disorder (OCD) using the Obsessive-Compulsive Inventory–Revised (OCI-R), social anxiety using the Liebowitz Social Anxiety Scale (LSAS), impulsivity using the Barratt Impulsiveness Scale (BIS-11), schizotypy using the Short Scales for Measuring Schizotypy (SSMS), alcohol misuse using the Alcohol Use Disorders Identification Test (AUDIT), eating attitudes using the Eating Attitudes Test (EAT-26), and apathy using the Apathy Evaluation Scale (AES). We also estimated cognitive ability with the ICAR test (16 items). In all histograms, the horizontal axis is bounded between minimum and maximum score values for each questionnaire. (**b**) Factor loadings for each questionnaire item on each symptom dimension: Anxious-Depression (AD, top), Compulsive Behavior and Intrusive Thoughts (CIT, middle), and Social Withdrawal (SW, bottom). The right panels display the distribution of scores on each dimension across participants. (**c**) Participant demographics. Distributions of gender (top) and age (bottom).

Bayesian model selection (see Methods) indicated that both sources contributed to decision variability across conditions (Fig. 2b; exceedance *p* > 0.999). Participants used higher learning rates in the V+ (volatile) condition (Fig. 2c; signed-rank test, V+ vs Ref: *z* = 9.8, *p* < 0.001), but also lower learning noise (V+ vs Ref: *z* = −3.1, *p* < 0.001) and increased choice temperature (V+ vs Ref: *z* = 4.4, *p* < 0.001). By contrast, these three parameters did not differ between the S+ (stochastic) and the reference condition. These selective differences between conditions are adaptive in the sense that the V+ condition requires faster learning and more frequent exploration of choice options due to the drifting values of choice options.

### Dimensional analysis of mental health symptoms

To study the relation between compulsivity and task behavior, we also administered a battery of nine self-report questionnaires assessing a large range of mental health symptoms known to fluctuate across the general population^6,8^ (first dataset: Fig. 3; second dataset: Supplementary Fig. 4). Participants’ mental health profiles varied within and across questionnaires, with most participants scoring within the normal range of symptoms (Fig. 3a).

**Figure 4.**
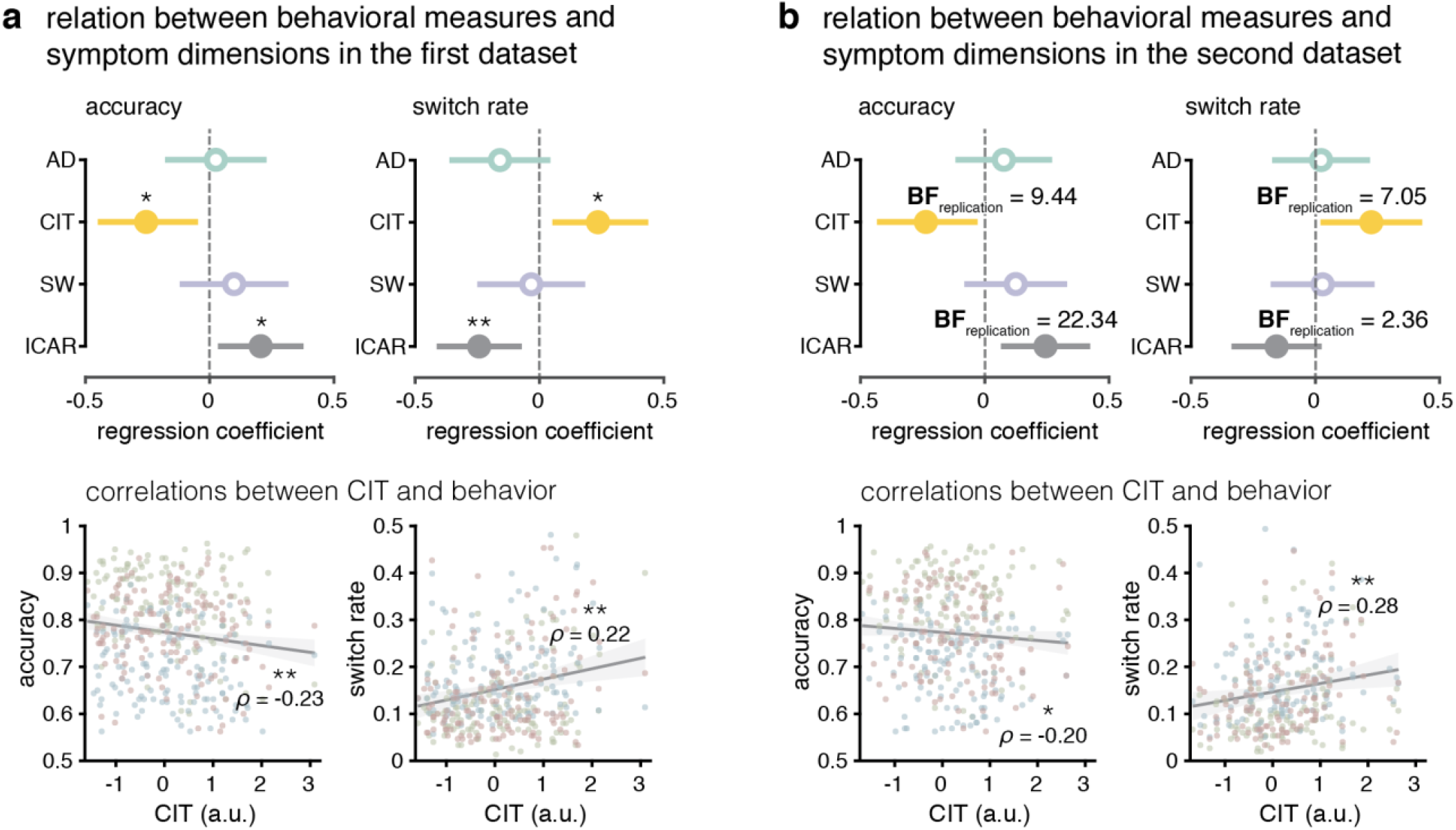
Relation between behavioral measures and symptom dimension scores. (**a**) First dataset: *N* = 137 participants. Top: regression analyses examining the associations between accuracy (left) and switch rate (right) as a function of AD, CIT, and SW scores along with a number of covariates among which cognitive ability (ICAR-16). Dots display regression coefficients and error bars are 95% CIs. \**p*<0.05, \*\**p*<0.01, statistical significance of regression coefficients. Bottom: correlation between CIT score and accuracy (left), and switch rate (right). Shaded area indicates the 95% CIs of the best fitting regression line. Circles indicate individual data points for each of the conditions (three data points per participant, green: Ref, red: S+, blue: V+). (**b**) Second dataset: *N* = 123 participants. Same analyses as in (**a**). Replication Bayes factor are shown for all effects that were significant in regression analyses in the first dataset.

Due to the strong correlations between questionnaire-specific scores, we employed a previously validated dimensional approach to the analysis of item-specific symptom scores^8^. Using a factor analysis (see Methods), we extracted three symptom dimensions from the covariance structure of responses to the 209 questionnaire items (Fig. 3b): an ‘anxious-depression’ (AD) dimension, a ‘compulsivity and intrusive thoughts’ (CIT) dimension, and a ‘social withdrawal’ (SW) dimension.

We also collected age and gender, along with a measure of general cognitive ability using the ICAR-16 test^18,19^, used as control variables in analyses of the relation between symptom dimensions and task behavior (Fig. 3c).

### Associations between symptom dimensions and decision variability

We first regressed accuracy and switch rate, averaged across task conditions, against the three symptom dimensions and control variables (Fig. 4a; see Methods). We found that accuracy was associated negatively with CIT (*β* = −0.255 ± 0.010, *p* = 0.011) and positively with ICAR-16 (*β* = 0.205 ± 0.088, *p* = 0.021). By contrast, switch rate was associated positively with CIT (*β* = 0.235 ± 0.099, *p* = 0.019) and negatively with ICAR-16 (*β* = −0.243 ± 0.087, *p* = 0.006). These results mean that CIT correlates negatively with accuracy (rank correlation, *ρ* = −0.230, *p* = 0.007) and positively with switch rate (*ρ* = 0.220, *p* = 0.010). These effects of CIT were replicated in the second dataset (Fig. 4b; accuracy: replication Bayes Factor (BF) = 9.44; switch rate: replication BF = 7.05). Neither accuracy nor switch rate was associated with AD or SW scores.

We then regressed learning noise and choice temperature against the three symptom dimensions and control variables (Fig. 5a). We found no association between learning noise and any symptom dimensions, but a positive association between choice temperature and CIT (*β* = 0.291 ± 0.096, *p* = 0.003) and a negative association between choice temperature and ICAR-16 (*β* = −0.202 ± 0.085, *p* = 0.019). This result means that CIT correlates positively with choice temperature (*ρ* = 0.276, *p* = 0.001), but not with learning noise (*ρ* = 0.108, *p* = 0.208). This selective association between CIT and choice temperature was replicated in the second dataset (Fig. 5b; replication BF = 9.8).

**Figure 5.**
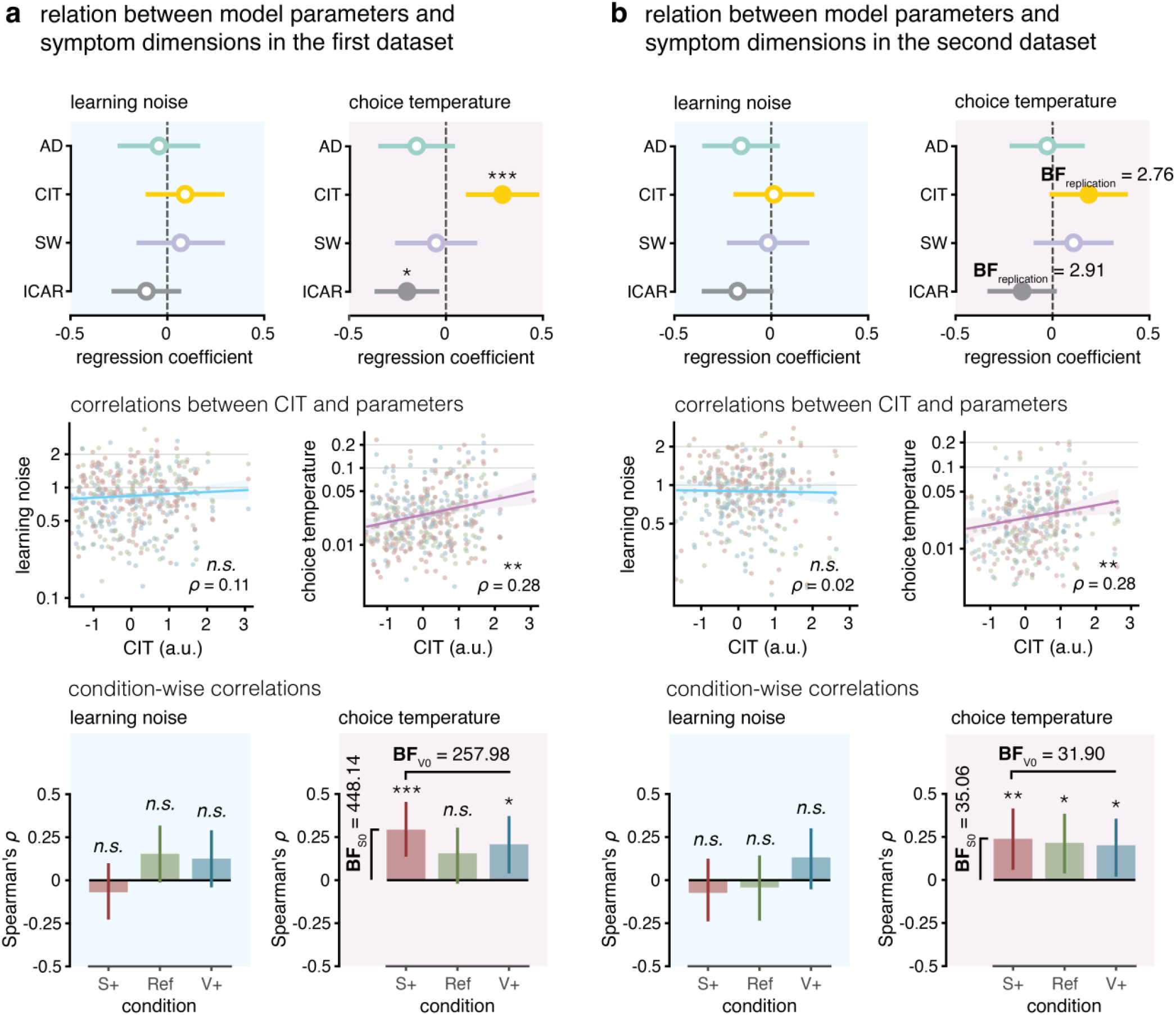
Relation between model parameters and symptom dimension scores. (**a**) First dataset: *N* = 137 participants. Top: regression analyses examining the associations between learning noise and choice temperature as a function of AD, CIT, and SW scores with a number of covariates among which cognitive ability (ICAR-16). Dots display regression coefficients and error bars are 95% CIs. \**p*<0.05, \*\**p*< 0.01, \*\*\**p*<0.001, statistical significance of regression coefficients. Middle: correlations between CIT score and learning noise (left), and choice temperature (right). Shaded area indicates the 95% CIs of the best-fitting regression line. Circles indicate individual data points for each of the conditions (three data points per participant, green: Ref, red: S+, blue: V+). Bottom: correlation between CIT and learning noise (left), and choice temperature (right), separately in each of the three conditions. Bar and error bars represent median Spearman coefficient and its 95% CI obtained using bootstrapping. \*\*\**p*<0.001, \**p*<0.05, *n*.*s*.: not significant. (**b**) Second dataset: *N* = 123 participants. Same analyses as in (**a**). Replication Bayes factors are shown for all effects that were significant in regression analyses in the first dataset.

As a control analysis, we regressed accuracy, switch rate and choice temperature against each of the nine questionnaire-specific scores (Supplementary Fig. 5; see Methods). Out of all questionnaires, we found that all these measures of variability were associated selectively with only the OCI-R score reflecting obsessive-compulsive symptoms^20^ (accuracy: *p* = 0.006; switch rate: *p* = 0.011; choice temperature: *p* = 0.001). Notably, no significant association was observed with the questionnaire-specific score measuring schizotypy in the general population^21^, despite the significant contribution of its items to the CIT dimension (Fig. 3b).

The positive association between CIT and choice temperature may either be adaptive if it is selective of the volatile condition where choice variability usefully drives exploration of choice options (account #1), or maladaptive if it is present even in the stochastic condition with stable option values (account #2). In line with account #2, we found that CIT correlated positively with choice temperature in both conditions (Fig. 5a; V+: *ρ* = 0.208, *p* = 0.015; S+: *ρ* = 0.293, *p* < 0.001). This pattern confers decisive evidence for a maladaptive account of this association (BF_V0_ = 258.0; see Methods). This maladaptive association between CIT and choice temperature in the stochastic condition was replicated in the second dataset (Fig. 5b; BF_V0_ = 31.9).

Although the positive association between CIT and choice temperature is maladaptive, we found no association between CIT and the magnitude of changes in learning rate, learning noise and choice temperature in the volatile condition (learning rate: *p* = 0.916, BF_1/0_ = 0.184; learning noise: *p* = 0.481, BF_1/0_ = 0.230; choice temperature: *p* = 0.748, BF_1/0_ = 0.192). This means that high CIT scores were associated with increased choice temperature, but not with a decreased adaptation of learning noise or choice temperature to volatile options.

### Trait-state and mediation analyses of decision variability

Despite its statistical significance, the positive association between CIT and choice temperature only explains a limited fraction of individual differences in choice temperature (explained variance = 8.4%). However, the meaning of this association strongly depends on two considerations. The first consideration is the trait-state structure of choice temperature. The CIT dimension, like the self-report questionnaires from which it is constructed, is thought to reflect compulsive behaviors in everyday life – a dominantly ‘trait’ measure. By contrast, choice temperature has been observed to vary at much faster time scales (e.g., between conditions in our task), and is therefore likely to include a substantial ‘state’ component. This state component decreases by nature the observed association between CIT and choice temperature.

To unpack individual differences in choice temperature into quantifiable trait and state components, we conducted a test-retest analysis in a subgroup of participants (*N* = 100 from the first dataset; see Methods). Accuracy, switch rate, learning noise and choice temperature all showed fair-to-good reliability (Fig. 6a; accuracy: intra-class correlation (ICC) = 0.441, *p* < 0.001; switch rate: ICC = 0.709, *p* < 0.001; learning noise: ICC = 0.468, *p* < 0.001; choice temperature: ICC = 0.718, *p* < 0.001). These results mean that the trait (i.e., test-retest reliable) component of learning noise explains about a fourth (22.2%) of its observed variance. In contrast, the trait component of choice temperature explains about half (52.0%) of its observed variance (Fig. 6b). Assuming that CIT reflects trait compulsivity and can therefore only explain the trait component of choice temperature, this means that CIT explains 8.4% of the overall variance in choice temperature, but a sixth (16.5%) of its trait component (Fig. 6b). By contrast, CIT explains less than 5% of the trait component of learning noise.

**Figure 6.**
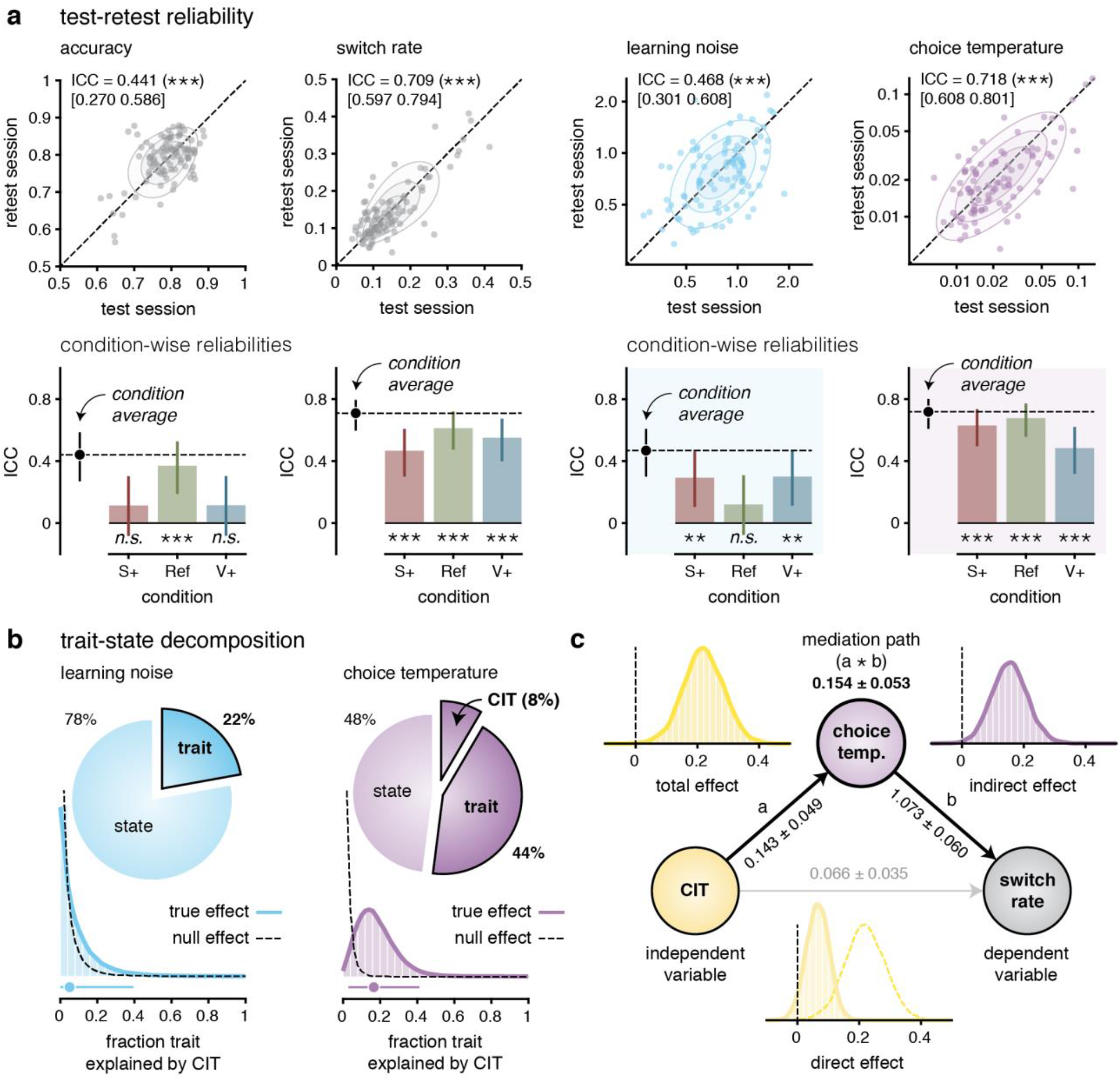
Trait-state and mediation analyses of decision variability. (**a**) Test-retest reliabilities of the two behavioral measures (accuracy, switch rate) and the two model parameters (learning noise, choice temperature) evaluated in the same participants using intraclass correlations (ICC) (*N* = 100 participants from the first dataset). Top: reliability of accuracy, switch rate, learning noise and choice temperature (averaged across conditions) between the test (x-axis) and retest (y-axis) sessions. Bottom: breakdown of ICCs for each of the three conditions (S+, Ref, V+). The black dot indicates the ICC for condition-averaged behavioral measures and model parameters. (**b**) Trait-state decomposition of learning noise and choice temperature. The trait component is defined as the variance in each model parameter that is shared across test and retest sessions. We assume that CIT is associated only with the trait component of each model parameter. (**c**) Mediation analysis comparing the indirect effect of CIT on switch rate through choice temperature (black paths a and b) against the direct effect of CIT on switch rate (grey path). Choice temperature significantly mediates the effect of CIT on switch rate.

The second consideration that constrains the significance of the association between CIT and choice temperature is the extent to which this association mediates the relation between CIT and observed behavioral variability (i.e., the fraction of switches between the two choice options). We reasoned that the association between CIT and choice temperature is only meaningful if it explains a significant fraction of the observed increase in switch rate with CIT. A mediation analysis^22,23^ (see Methods) confirmed this prediction. Indeed, we found a significant mediation effect of choice temperature on the relation between CIT and switch rate (Fig. 6c; *β* = 0.154 ± 0.053, *p* = 0.004), together with a weak direct effect of CIT on switch rate (*β* = 0.066 ± 0.035, *p* = 0.066) – down from a strong total effect (*β* = 0.219 ± 0.062, *p* < 0.001).

## Discussion

Compulsivity has been linked to a broad range of shifts in decision-making under uncertainty^1^. Compulsive individuals typically show indecisiveness when decision commitment can be postponed^3,5^, but also increased decision variability^4^ and impoverished decision policies in hierarchical environments^10,11^. These effects may arise from shifts in how compulsive individuals choose between uncertain options. However, these effects may also result from upstream changes in the learning of option values from variable outcomes. Here, by decomposing behavioral variability into its learning and choice components, we found that under uncertainty, compulsive individuals make variable and maladaptive choices, but do not learn the values of choice options less precisely than others.

Our findings confirm that compulsive individuals make less accurate decisions and switch more often between uncertain choice options than other individuals. However, we show that compulsive individuals do not learn the values of these choice options less precisely than others. Rather, they differ from others in that they choose more variably than others – a finding which extends recent reports by accounting for the presence of learning imprecisions^3,24^. This selective association between compulsivity and choice stochasticity is consistent with the observation that compulsivity is more strongly related to individual differences in executive than cognitive parameters^6^. Indeed, the stochasticity of participants’ choices is arguably more dependent on executive control^25^ than on the learning and representation of option values during reinforcement learning. This pattern of findings is also in line with previous work showing that clinically-diagnosed OCD patients have difficulty with decision-making despite holding accurate representations of task variables and intact confidence estimates^4^.

Using a test-retest approach, we made use of the empirical reliability of model parameters to decompose learning noise and choice temperature into state and trait components. This analysis showed that individual differences in compulsivity alone explain about a sixth (16.5%) of the trait variance in choice temperature. This quantitative result is important, because most previous studies in computational psychiatry have (like us) attempted to relate personality or psychiatric traits – typically evaluated via self-report questionnaires – to model parameters measured in a specific task, such as choice temperature in our study. However, most of these studies typically do not measure the test-retest reliability of model parameters, and therefore typically assume a degree of stability of these parameters across time. This ‘trait’ component of model parameters, and task-specific variables in general, is often much lower than the ‘trait’ component of self-report questionnaire-specific scores^26,27^. Indeed, most behavioral tasks in cognitive science focus on the within-subject comparison of different experimental conditions^28^. Quantifying the specific trait association between psychiatric symptoms and model parameters has critical implications for cognitive phenotyping and, in the longer term, for the possible identification of computational profiles for personalized psychiatry.

Here, we re-tested the same individuals in the same task conditions to decompose choice variability into state and trait components, in line with previous work which has quantified the trait and state components of risk-seeking behavior, or general intelligence^29,30^. For instance, preference for information seeking has been shown to be relatively stable^24^, like certain aspects of preference for costly information, even in conditions of decreased immediate expected reward^31^. Decision parameters in general also show large interindividual differences that are relatively stable over time^32^. Here, we established that choice temperature (reflecting the stochasticity of the choice policy) has the largest trait component of all parameters of our reinforcement learning model (52% of its overall variance). The other source of decision variability, arising from learning imprecisions, has a lower but still significant trait component (22% of its overall variance). It is important to note that having a higher trait component is not inherently ‘better’ than having a lower trait component. Indeed, the respective variances of state and trait components may reflect a flexibility-consistency trade-off in model parameters. For example, the state component may reflect online, moment-to-moment tuning of model parameters – as a function of available attentional/cognitive resources and environmental demands, to allow for adaptive behavior across changing task conditions. The fact that both learning noise and choice temperature differ between conditions in our study confirms that the state component does not reflect random noise in model parameters but rather an adaptation to specific task conditions.

Interestingly, we did not find any significant association between learning noise and any symptom dimension, despite its substantial interindividual differences and its significant (albeit moderate) trait component. Which personality or psychiatric trait(s) may explain individual differences in learning noise therefore remains to be identified. One possibility is that learning noise is a highly state-dependent, flexible cognitive parameter for which within-individual variations across tasks and contexts are much larger than between-individual variations. Given that learning noise (and more generally internal noise in cognitive computations) is an important source of decision variability under uncertainty^12,13,33^, examining its trait-state structure and its relation to personality and psychopathology constitutes a major avenue for future research.

Our results show that compulsivity is associated with increased choice variability across task conditions, irrespective of whether this type of ‘random’ exploration is adaptive (when option values drift over time) or maladaptive (when option values are stable). This finding may potentially inform the development or refinement of cognitive behavioral therapies (CBTs), which aim at identifying which cognitive processes should be targeted to improve behavioral symptoms^34^. By revealing a selective impact of compulsivity on choice variability without alterations in reinforcement learning, our study highlights executive control – rather than learning itself – as a possible target for treatment. Future work should validate this finding, obtained using subclinical differences in self-reported compulsivity in the general population, in patients diagnosed with OCD^4,25^.

## Methods

### Experimental protocol

Our experimental design comprises three conditions, each divided into two-armed bandit tasks^17^. Rewards were drawn between 1 and 99 points. The Reference (Ref) and Stochastic (S+) conditions had fixed reward distributions. In contrast, in the Volatile (V+) condition the means of the reward distributions associated with each arm followed a random walk over time. In all conditions, the mean rewards associated with each option were mirrored relative to 50 points. In the V+ condition, we define a reversal as the moment where the mean rewards associated with each of the two options crossed each other. Participants underwent two blocks of 80 trials of each condition.

The two bandits were displayed as colored discs for the V+ condition (Fig. 1). By contrast, the two bandits were displayed as black shapes for the Ref and S+ conditions. In each new round, new shapes were presented, so as to avoid carry-over effects across rounds and ensure that participants reset and treat each new round separately from the previous one. We created these displays to ensure participants distinguish between stable (Ref and S+) vs. volatile (V+) conditions. After each decision, the chosen option was briefly displayed in the center of the screen, and followed by the points obtained on this trial (Fig. 1a). Upon completion of the task, participants earned £3.30, plus an additional £1.00 bonus if their accuracy was significantly better than chance level throughout the whole experiment.

Critically, we sought to achieve matched task difficulty between the S+ and V+ conditions. To achieve this, we used simulations from an optimal agent playing the task (see also Computational reinforcement learning model). For each participant, 2,000 random walks were generated with rewards drawn from a beta distribution with initial mean of 50 points and drifting standard deviation of 5 points. We subsequently discarded all walks that had steps larger than 10 points or travelled below 10 or above 90 points. To create blocks of trials for the V+ condition, we selected excursions below or above 50 points of 8, 12, 16, 20 and 24 steps. We employed the mean reward for each excursion as the static mean reward of a round of trials for the Ref condition. Furthermore, we set the variance of sampled rewards so that 75% of rewards from the better option were higher than 50 points in the Ref condition. Then, we computed the effective sampling variance and drifting variance of rewards; we simulated a Kalman filter with greedy decisions to obtain its accuracy in the V+ condition. Finally, we generated the S+ condition by incrementally regressing the static mean reward from the Ref condition towards 50, until the accuracy of the Kalman filter in this S+ condition matched that of the V+ condition.

After providing informed consent, participants had to be in full-screen mode to be able to carry on. A set of instructions introduced participants to the task mechanics (e.g., response keys). They were instructed about the three experimental conditions that were signaled via written prompts on the screen. The Ref condition was always performed first. Then, the order of S+ and V+ conditions was counterbalanced across participants. Before each condition, a practice block was provided so participants get familiar with the condition. After each practice block, we provided them with a debriefing about their performance on the condition, with an illustration of the trajectory of their decisions and rewards throughout the block. They were offered breaks in-between blocks. The experiment was self-paced, but had to be completed within 60 minutes.

### Self-report questionnaires assessing mental health symptoms

After completing the task, participants completed a set of nine questionnaires. We sought to sample variability in mental health symptoms naturally fluctuating in the general population. We evaluated depression using the Self-Rating Depression Scale (SDS)^35^, trait anxiety using the State Trait Anxiety Inventory (STAI) Form Y-2, obsessive-compulsive disorder (OCD) using the Obsessive-Compulsive Inventory – Revised (OCI-R)^20^, social anxiety using the Liebowitz Social Anxiety Scale (LSAS)^36^, impulsivity using the Barratt Impulsiveness Scale (BIS-11)^37^, schizotypy using the Short Scales for Measuring Schizotypy (SSMS)^21^, alcohol misuse using the Alcohol Use Disorders Identification Test (AUDIT), eating attitudes using the Eating Attitudes Test (EAT-26), and apathy using the Apathy Evaluation Scale (AES) (Fig. 3). We also estimated cognitive ability with the short International Cognitive Ability Resource (ICAR) test^18,19^ (16 items), which we used as a covariate alongside gender and age in subsequent regression analyses. In addition to the task bonus, participants who completed the questionnaire battery earned an extra £3.00.

### Participants

Three groups of 200 adult participants were initially recruited via the Prolific Academic platform (prolific.co). Here, we only use two of these datasets. The third one was not usable because ICAR scores were substantially (and significantly) lower than their expected distribution at the level of the group, thereby compromising the quality of questionnaire ratings. Sample size was determined a priori such that a between-participant correlation of 0.20 could be detected at standard levels of 80% power and 0.05 of significance threshold. Participants provided informed consent through a digital form regarding their participation in the study, as approved by the ethical review committee of the Institut National de la Santé et de la Recherche Médicale (IRB #00003888).

Exclusion criteria allowed us to maintain both task and questionnaire data quality. First, participants whose task accuracy did not exceed chance level were excluded (one-tailed threshold of 0.05 in any of the three conditions (binomial test against 50% accuracy)). This step led to a remaining *N* = 154 participants (mean age = 29.6 ± 10.0 years, 53 females) in the first dataset, and *N* = 142 participants (mean age = 25.4 ± 6.4 years, 72 females) in the second dataset. Second, we excluded participants who missed a catch item embedded into one of the questionnaires. The final sample comprised *N* = 137 participants (Fig. 3) for the first dataset, and *N* = 123 for the second dataset. To quantify the test-retest reliability of our behavioral variables and model parameters, out of the first dataset, *N* = 100 participants completed a second time the experiment after an average of two weeks from their first participation.

### Transdiagnostic factor analysis

To address the heterogeneity within-questionnaire and the correlations between questionnaires, we performed a factor analysis with an oblique rotation on the 209 questionnaire item data of 497 participants from our previous study^6^. The factor weights obtained were then applied on the questionnaire responses in our first and second datasets separately. Given the item-participant ratio here, instead of performing a de novo factor analysis, we leveraged the larger sample size obtained in our previous study. The factor analysis indicated that a three-factor solution best and most parsimoniously explained the data. In line with previous work^6,8^, these ‘transdiagnostic’ factors were labelled on the basis of questionnaire items loadings most strongly on each of these factors: ‘Anxious-Depression’ (AD), ‘Compulsive Behavior and Intrusive Thought’ (CIT), and ‘Social Withdrawal’ (SW). Consistent with previous work, we observed that AD and CIT factors were positively correlated (*ρ* = 0.357; *p* < 0.001).

### Computational reinforcement learning model

#### Model description

After each reward, the posterior value of the chosen option 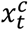 is updated using a Kalman filter corrupted by random additive noise *ε*_*t*_. *ε*_*t*_ is drawn from a Gaussian distribution with a mean of zero and a standard deviation *η*_*t*_:

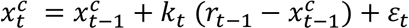

where *k*_*t*_ is the Kalman gain. It depends on the posterior variance of the chosen option 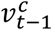 and the sampling variance *υ*_*s*_ of outcomes. The latter was set to its true effective value (0.0163 for outcomes rescaled between 0 and 1) and used as a scaling parameter:

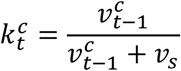

We updated the posterior variances of the two choice options *o* ∈ {1,2} with the standard equation of the Kalman filter:

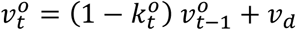

where *υ*_*d*_ is the drifting variance assumed by the Kalman filter. *υ*_*d*_ was estimated in terms of its associated asymptotic Kalman gain *α* for an option that would be selected on *n* → ∞ trials. In line with recent results^13^, the standard deviation of the random noise corrupting the update of the posterior value of the chosen option scaled as a constant fraction *ζ* of the update 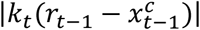. We initialized the posterior variances of the two options to the true variance of mean outcome values across all conditions (0.0214 for outcomes rescaled between 0 and 1).

The posterior value of the unchosen option 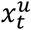 regresses toward the overall average reward of 50 points across the two options (or 0.5 for rewards rescaled between 0 and 1). This regression-to-the-mean is controlled by an exponential decay factor *δ*:

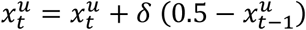

The probability of choosing option 1 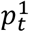 based on the posterior values of the two options then follows a standard softmax policy with choice temperature *τ*:

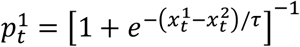

#### Model fitting

We applied Sequential Monte Carlo methods to estimate the conditional likelihoods of the decisions of each participant given the set of parameter values. We employed Bayesian Adaptive Direct Search (BADS) with 10 random starting points for each parameter *α, ζ, δ* and *τ* ^38^ to first obtain point estimates of best-fitting parameter values. We then used these estimates as starting points for the estimation of their joint posterior distribution using Variational Bayes Monte Carlo (VBMC) ^39,40^.

#### Model comparison

We performed a model comparison to examine whether the addition of learning noise (controlled by its Weber fraction *ζ*) and soft choices (controlled by their temperature *τ*) were both necessary to fit participants’ decisions. We ran a random-effects Bayesian Model Selection (BMS) using its standard Dirichlet parameterization d ^41^. As in previous research ^13^, we compared the model including both sources of decision variability to two model variants. A first model variant had no learning noise (*ζ* = 0). A second model variant had a greedy choice policy with *τ* = 0. The full model outperformed the other two model variants (model prevalence of 70% for the full model, exceedance *p* > 0.999) (Fig. 2b).

#### Model recovery

We performed model recovery to ensure that the models we would compare were both differentiable and recoverable in the fitting procedure. We simulated the model from the parameters obtained in the comparison procedure, and fit all three models on this simulated data. and performed model comparison for each of these three fits. The model prevalence and their protected exceedance probabilities showed each model as distinguishable and recoverable (see Supplementary Figure 3a).

#### Parameter recovery

We performed parameter recovery analyses to ensure that there were no systematic relationships between the parameters generated by our model-fitting procedure. First, we destroyed any correlation between the best-fitting parameters via random permutation, and generated simulated data from these parameter sets. Then we fitted the model to this simulated data and calculated the resultant confusion matrix (Supplementary Figure 3b). The lack of significant correlations in the off-diagonal verifies that our fitting procedure does not generate any systematic correlations between model parameters. We then confirmed that the model parameter fits to participants’ data are themselves recoverable via simulation and fitting of the generative model using these parameter values (Supplementary Figure 3c).

### Statistical analyses

We used Wilcoxon signed-rank tests for comparing (i) model-free variables, accuracy and switch rate; and (ii) best-fitting model parameters between conditions. We employed non-parametric Spearman correlations unless specified otherwise. We computed standard errors on Spearman’s rank correlation coefficients (Fig. 5) using bootstrapping. We randomly sampled datasets from the original dataset (with identical sample size and replacement) 1,000 times and calculated the correlation on these new datasets.

To investigate the associations between mental health symptoms and task behavior, we conducted regression analyses (Fig. 4 and 5). We specified accuracy, switch rate, and model parameters in turn as dependent variables, in separate regressions. Predictors were scores on the three symptom factors, alongside age, gender and cognitive ability (ICAR) as additional control variables. Statistical significance was assessed using one-sample t-tests on regression coefficients. To directly illustrate the associations between task variables and symptom scores identified, we also examined Spearman correlations between CIT and accuracy, switch rate (Fig. 4), learning noise, and choice temperature (Fig. 5). To examine whether the identified associations replicated in our second dataset, we computed Bayes Factor (BF) tests for replication success^42^ answering the question “Is the effect from the replication attempt comparable to what was found before, or is it absent?”.

To examine the maladaptation of choice temperature in the stochastic condition (S+) with stable option values, we computed two Bayes Factors tests on the correlation coefficients between choice temperature and CIT. First, we calculated the relative evidence of finding a zero correlation in a null distribution over finding it in the distribution of S+ (Fig. 5; first dataset: BF_S0_ = 448.1; second dataset: BF_S0_ = 35.1). We then calculated the relative evidence of finding the true S+ correlation from the distribution of V+ over finding it in a null distribution (Fig. 5; first dataset: BF_V0_ = 258.0; second dataset: BF_V0_ = 31.9). These Bayes Factor tests were obtained by calculating the likelihood ratios of the marginal probability density functions derived from normal distributions generated by values of Fisher *z*-transformations on Spearman’s correlation coefficients.

To examine reliability between test and retest, we calculated intraclass correlations (ICC) for accuracy, switch rate, and model parameters measured at test and retest sessions in the same participants (Fig. 6a). We employed a two-way mixed-effects, absolute agreement model on single measurements used in test-retest reliability studies^43^. All variables were log-transformed before calculating ICC scores. Statistical significance was assessed via *F*-tests. Furthermore, we performed a mediation analysis^22,23^ to examine whether choice temperature mediates the link between CIT and the measured behavioral variability (Fig 6b). We reasoned that the link between CIT and choice temperature only matters if it explains to some degree the increase in switch rate with increased CIT score. Hence, two pathways were specified. A direct pathway from CIT to switch rate was compared against an indirect pathway from CIT to choice temperature to switch rate.

### Data and code availability statement

Anonymized behavioral and questionnaire data, together with MATLAB code for model fitting and regression-based analyses of individual differences in task behavior, model parameters and mental health symptoms, are available on a public repository: https://github.com/jlexternal/RLVOLUNP_CIT_ana

## Author contributions

Conceptualization: VW. Methodology: JKL, MR, VW. Investigation: JKL. Visualization: JKL, VW. Supervision: VW. Writing—original draft: JKL, MR, VW. Writing—review & editing: JKL, MR, VW.

## Acknowledgements

This work was supported by a starting grant from the European Research Council (ERC-StG759341) awarded to V.W., and by a department-wide grant from the Agence Nationale de la Recherche (ANR-17-EURE-0017, EUR FrontCog). M.R. is the beneficiary of a postdoctoral fellowship from the AXA Research Fund, and is supported by La Fondation des Treilles. The funders had no role in study design, data collection and analysis, decision to publish or preparation of the manuscript.

