## Supplementary Figures for "Compulsivity is linked to maladaptive choice variability but unaltered reinforcement learning under uncertainty"

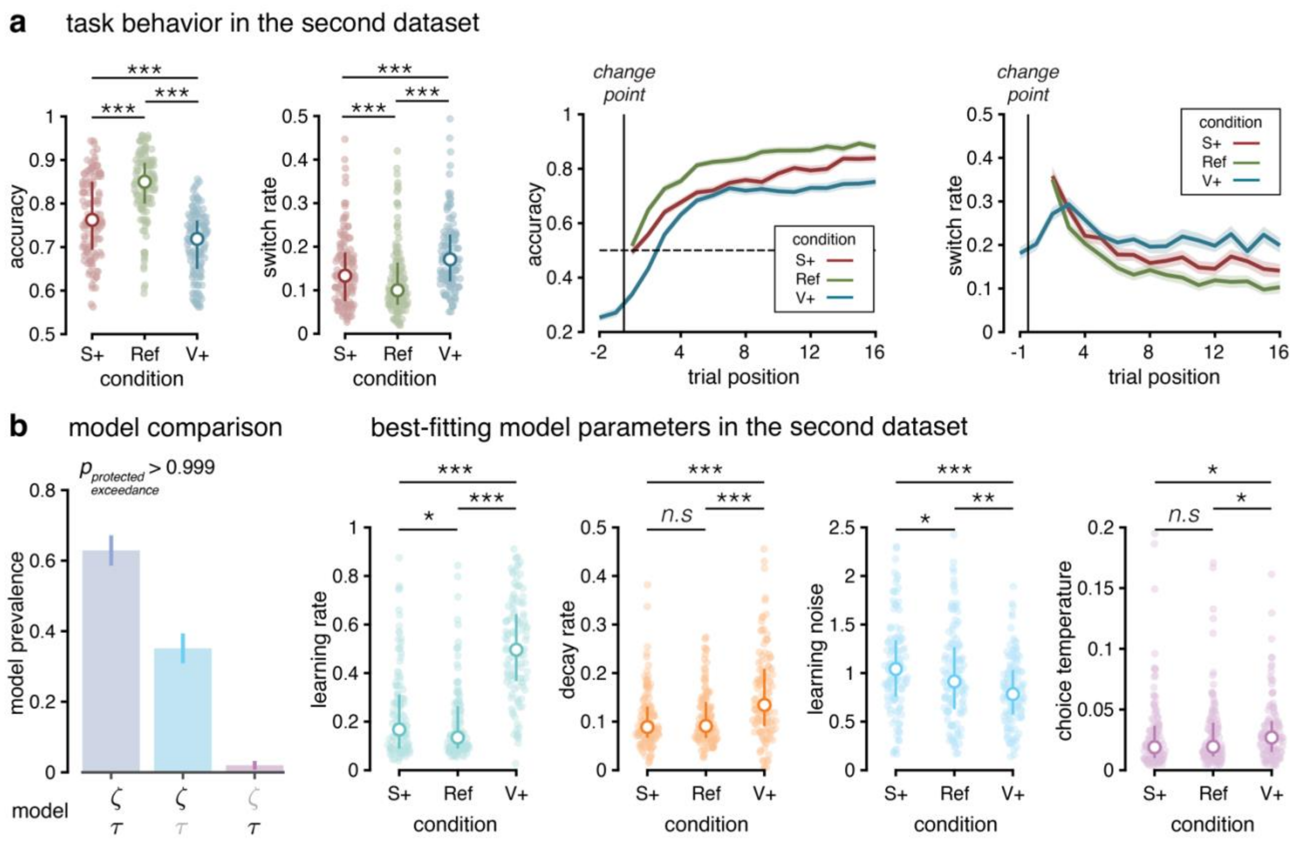


**Supplementary Figure 1. Participant behavior and modeling results in the second dataset** (second dataset: $N = 123$) (**a**) Left: Average accuracy and average switch rate in each condition. Colored dots represent individual participants' mean accuracy and overall switch rate respectively ($N = 137$). White dots indicate the median accuracy and median switch rate respectively. Error bars represent the 1st and 3rd quartiles. Right: Accuracy and proportion of switch decisions as a function of trial position in each condition. Solid lines indicate the mean accuracy and mean switch rate across participants. The vertical line represents the start of a new block (Ref, S+) or a reversal (V+). (**b**) Left: Model comparison favors a model with both learning noise and choice stochasticity (left bar), with high prevalence across tested participants. Models lacking either source of decision variability (middle and right bars) provide poorer fits to human behavior. Right: Best-fitting individual model parameters in each of the three conditions. White dots indicate the median parameter value and error bars represent the 1st and 3rd quartiles. Circles display individual data points. ***$p<0.001$, **$p<0.01$, *$p<0.05$, *n.s.*: not significant, signed-rank tests.

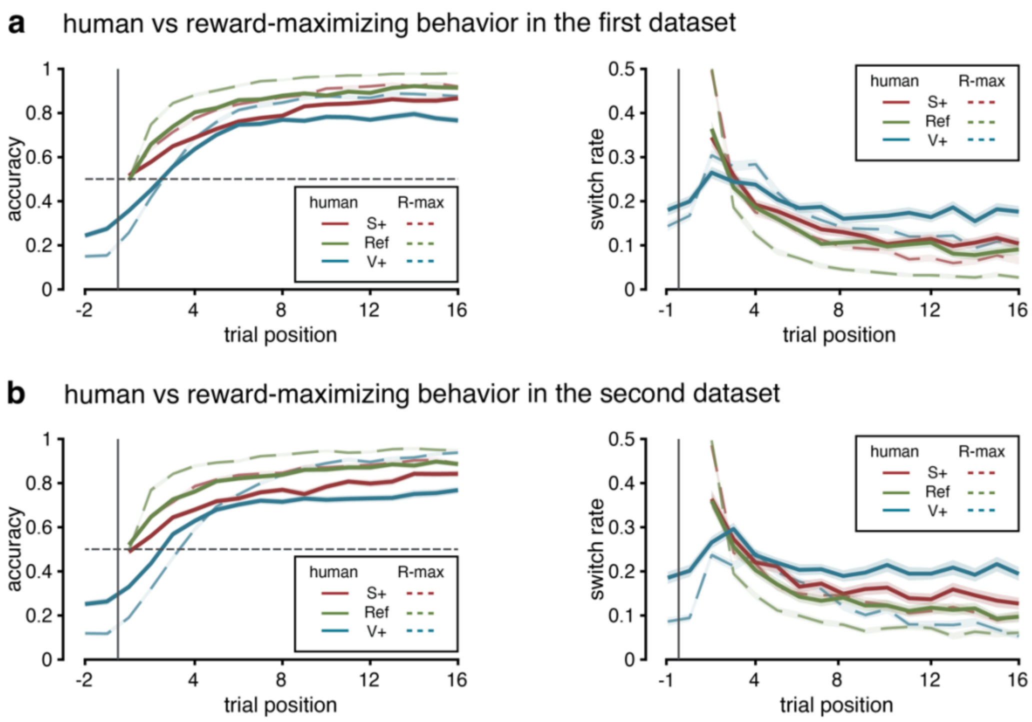


**Supplementary Figure 2. Human vs reward-maximizing model simulation behavior.** (**a**) Accuracy and proportion of switch decisions as a function of trial position in each condition in the first dataset ($N$ = 137). Solid lines indicate the mean accuracy and mean switch rate across participants. Dashed lines indicate the mean accuracy and mean switch rate across model simulations generated from reward-maximizing parameters. The vertical line represents the start of a new block (Ref, S+) or a reversal (V+). (**b**) Second dataset: $N$ = 123. Same analyses as in (**a**).

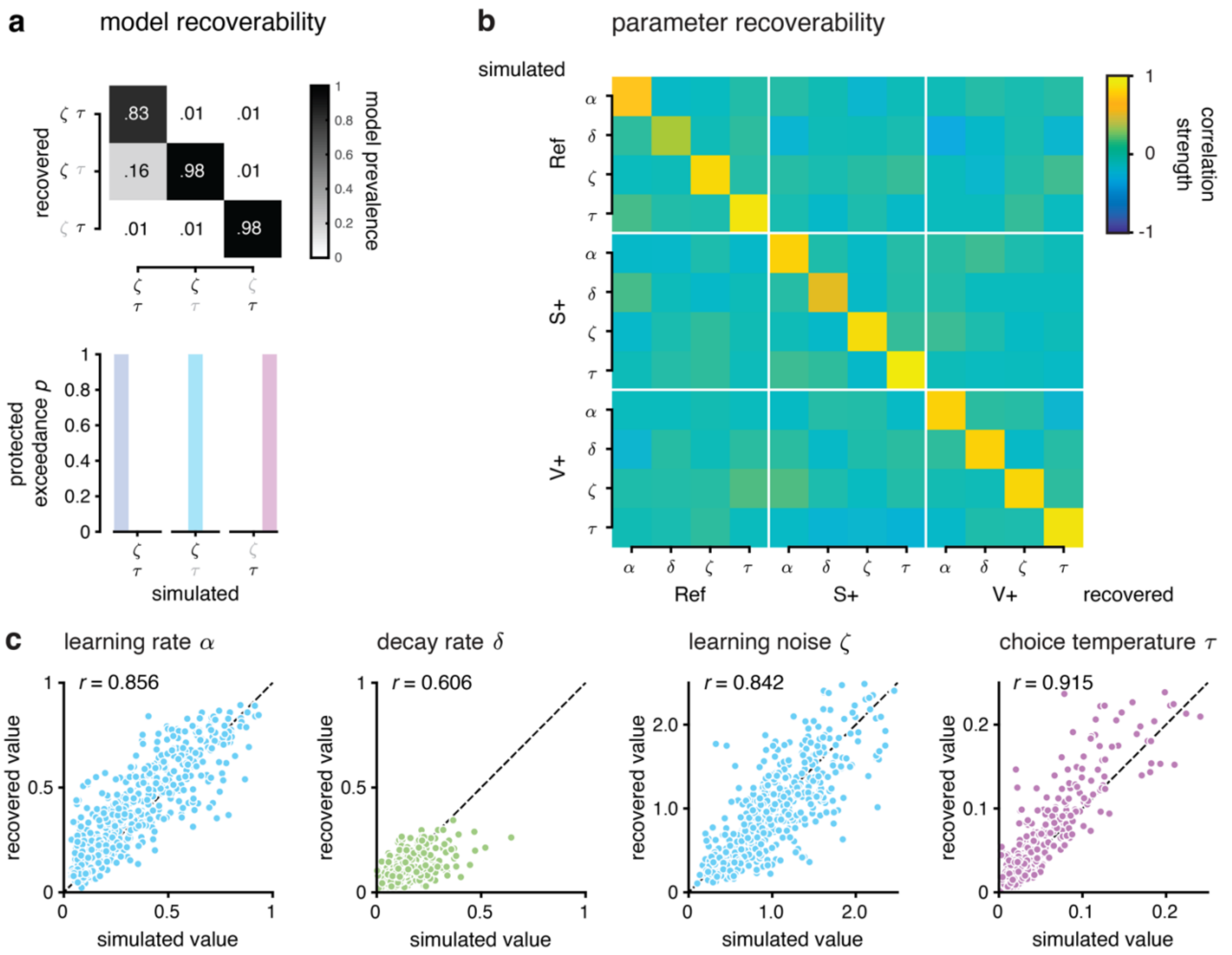


**Supplementary Figure 3. Model validation and parameter recovery analyses.** (**a**) Top: Model recovery matrix. Frequencies of recovered model on simulated data generated from three candidate models fitted on participants. Bold letters indicate inclusion of the parameter for learning noise or choice stochasticity. Bottom: Protected exceedance probabilities showing the prevalence of each generative model when compared with the others. (**b**) Parameter confusion matrix. Correlation matrix of simulated on shuffled parameter values and the resulting recovered model parameters. The near unit matrix provides evidence that any correlations in the true correlation matrix is not due to some systematic error. (**c**) Scatter plots of simulated and recovered parameter values.

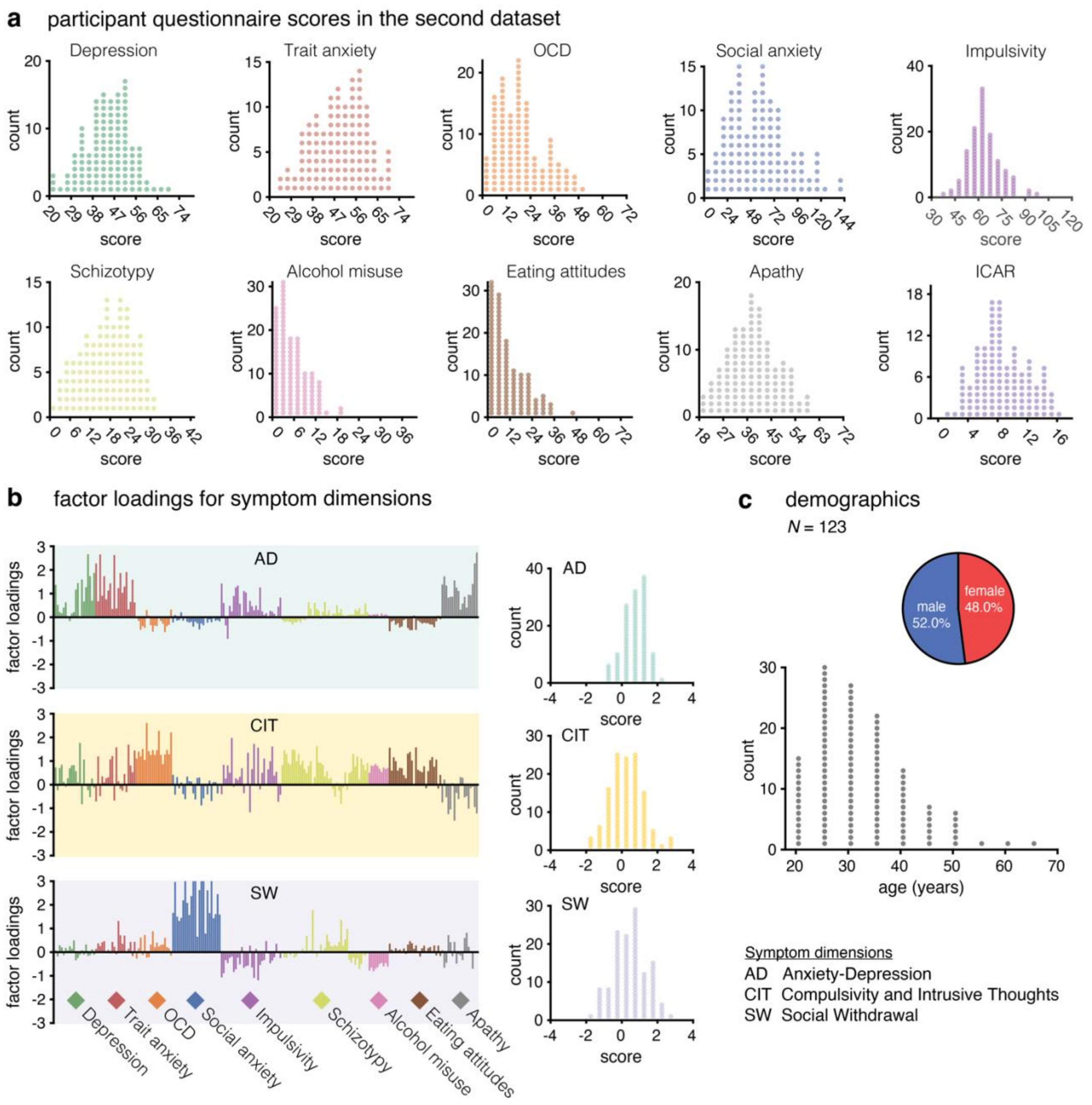


**Supplementary Figure 4. Inter-individual variability in symptom dimension scores and demographics variables** (Second dataset: *N* = 123 participants). (**a**) Histograms showing distributions across questionnaire scores in our sample. We evaluated Depression using the Self-Rating Depression Scale (SDS), Trait Anxiety questionnaire using the State Trait Anxiety Inventory (STAI) Form Y-2, Obsessive Compulsive Disorder (OCD) using the Obsessive-Compulsive Inventory–Revised (OCI-R), Social anxiety using the Liebowitz Social Anxiety Scale (LSAS), Impulsivity using the Barratt Impulsiveness Scale (BIS-11), Schizotypy using the Short Scales for Measuring Schizotypy (SSMS), Alcohol misuse using the Alcohol Use Disorders Identification Test (AUDIT), Eating attitudes using the Eating Attitudes Test (EAT-26), and Apathy using the Apathy Evaluation Scale (AES). We also estimated cognitive ability with the ICAR test (16 items). In all histograms, the horizontal axis is bounded between minimum and maximum score values for each questionnaire. (**b**) Factor loadings for each questionnaire item on each symptom dimension: Anxious-Depression (AD, top), Compulsive Behavior and Intrusive Thoughts (CIT, middle), and Social Withdrawal (SW, bottom). The right panels display the distribution of scores on each dimension across participants. (**c**) Participant demographics. Distributions of gender (top) and age (bottom) in the second dataset.

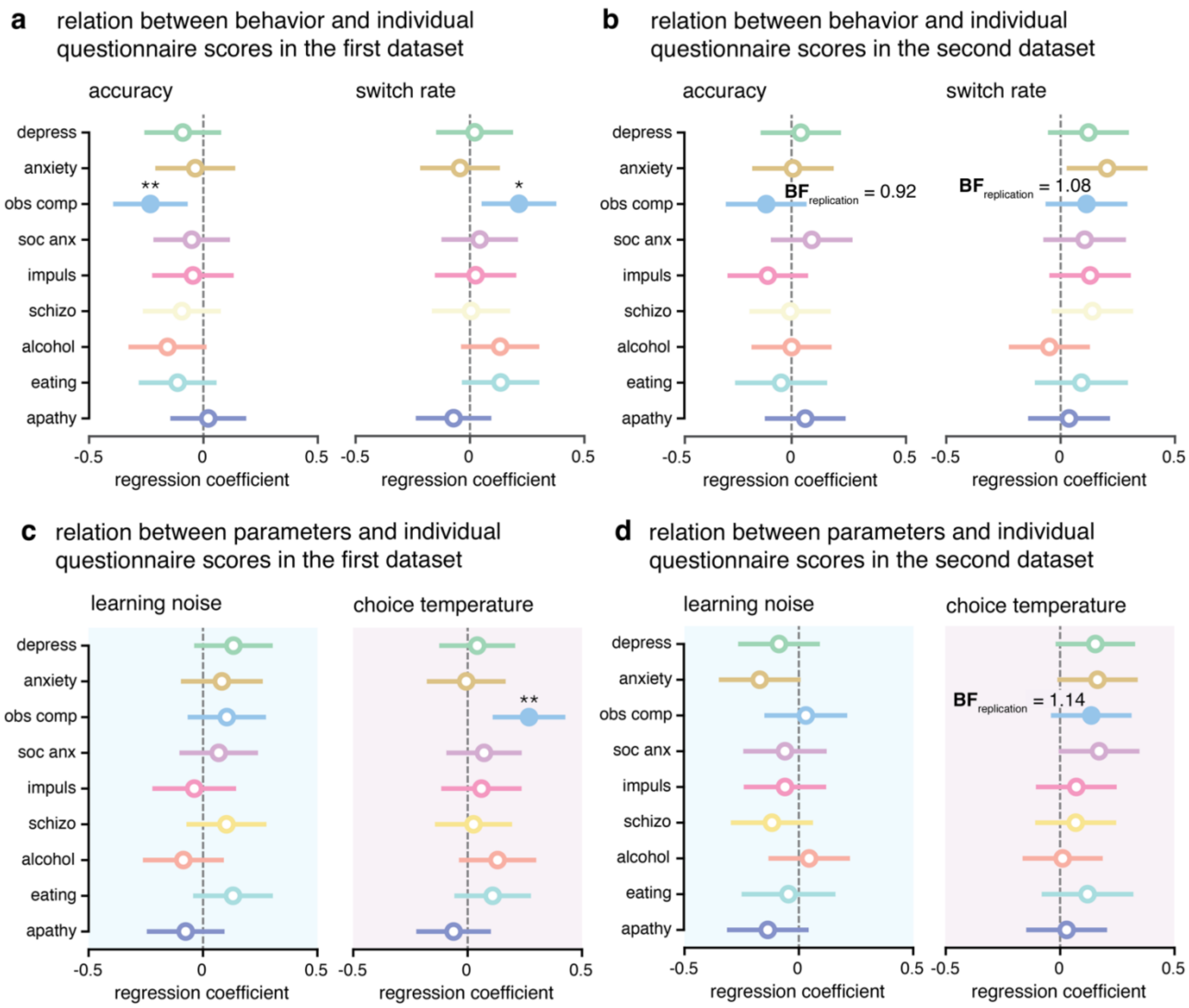


**Supplementary Figure 5. Relation between behavior and parameters with questionnaire scores.** (**a**) First dataset: $N$ = 137. Regression analyses examining the associations between accuracy (left) and switch rate (right) as a function of the individual questionnaire score totals. Dots display regression coefficients and error bars are 95% CIs. *$p$<0.05, **$p$<0.01, statistical significance of regression coefficients. (**b**) Second dataset: $N$ = 123. Same analyses as in (**a**). Replication Bayes factor shown wherever effects were significant in the first dataset. (**c**) First dataset. Regression analyses examining the associations between learning noise and choice temperature as a function of the individual questionnaire score totals. Dots display regression coefficients and error bars are 95% CIs. **$p$< 0.01, statistical significance of regression coefficients. (**d**) Second dataset. Same analyses as in (**c**). Replication Bayes factor shown for the second dataset wherever effects were significant in the first dataset.
